# On the interaction of replication and transcriptional regulation

**DOI:** 10.1101/2023.03.20.533469

**Authors:** Pietro Gemo, Matteo Brilli

## Abstract

DNA replication introduces a gradient of gene copy numbers, and in Bacteria it affects gene expression accordingly. In *E. coli* and other species, the slope of the gradient averaged over the population can be predicted on the basis of its relationship with growth rate. In this work we integrated this growth- and position-dependent gradient into a classical transcriptional regulation model, to highlight their interaction. The theoretical treatment of our model highlights that the sensitivity to transcription factor-mediated regulations acquires an additional dimension related to the position of a locus on the oriter axis and to division time. This reinforces the idea of replication as an additional layer in gene regulation. We highlight here that replication- and transcription factor-mediated regulations can in theory work in concert or counteract each other, and we discuss why this is important from an evolutionary point of view with respect to both steady state transcript abundance and its variance across conditions. Finally, we note that this treatment may improve the estimation of kinetic parameters for transcription factor activity using RNA-seq data, and the estimation of the dispersion factor in differential gene expression analysis when division time across conditions changes significantly.

## Introduction

Most Bacteria in the environment experience a so-called *feast-famine* cycle [15]: short periods of exponential growth in nutrient-rich conditions, followed by nutrient depletion and then by a starvation period. During the latter most cells die while the remaining slow down their growth rate significantly. Therefore, species that can grow faster in rich conditions can outcompete other bacteria in the community by consuming a much larger fraction of the available nutrients before another famine period [12]. *E. coli* implements a successful strategy in this sense: it is mono-ploid when slow growing - e.g. at *µ ≈* 0.7*h*^*−*1^ (*τ ≈* 1*h*) but in rich conditions, it can shift to a mero-oligoploid growth *mode*, whereby cells are able to fire multiple replication forks during the same cell cycle round (e.g. at *τ ≈* 25*min* or *µ ≈* 1.7*h*^*−*1^, and Figure 1) [9]. This growth *mode* enables *E. coli* and other species to divide faster than the time required to copy the chromosome, which in *E. coli* is relatively constant (*C ≈* 40*min*) [5]. While mono-ploid species have at most two copies of the same gene as a consequence of replication, mero-oligoploids can have up to 8 *−* 10 copies of a same locus (Figure 1).

**Fig. 1.**
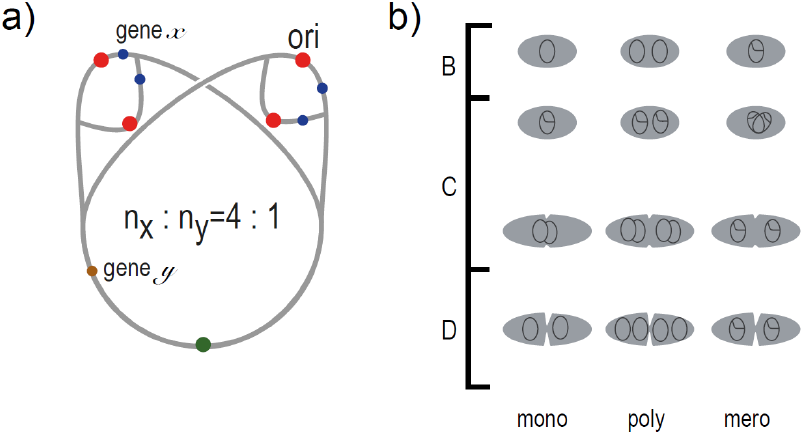
a) Representation of a circular chromosome with three active replication forks (*n*_*Ori*_ = 4). Genes *x* and *y*, are in a 4 : 1 ratio but this ratio changes with the number of replication forks. b) Cell cycle for mono-(exact), poly- and mero-oligoploid (idealized) species. *C* is the time needed for the replication of an entire chromosome. *D* is the time needed for segregation and cell division. In *E. coli B* is a *virtual* cell cycle phase, since during exponential phase newborn cells immediately start replicating the genome.

Copy numbers were shown to obey a relatively simple formula, defined since the late 60s [3, 5, 10]; in this framework, the copy number *n*_*a*_ of locus *a* located at position *p*_*a*_ on the Ori/Ter axis, can be calculated as:

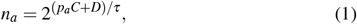

where *C, D* have the meaning as in Figure 1b and *τ* is the division time (*τ* = *ln*2*/µ*, with *µ* the growth rate); *p*_*a*_ is the relative distance of the locus from terminus such that *p*_*Ori*_ = 1 and *p*_*Ter*_ = 0. By introducing these parameters into Eq. 1, we get:

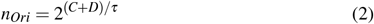

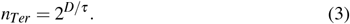

Although the model was based on *E. coli*, where it has an extremely good fit over a wide range of growth rates, it is similarly valid in other species [17]. Moreover, since Eq. 2 and 3 imply:

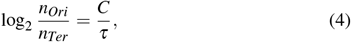

one can extrapolate the growth rate *in vivo* by exploiting shotgun metagenomic sequencing coverage. Indeed, the argument of the logarithm in Eq. 4 corresponds to the so-called peak-to-trough ratio (PTR) that was shown to be a good predictor of growth rate *in vivo* in different organisms [13, 18, 22, 11].

In Fondi et al., 2023 (doi: 10.1101/2022.09.05.506644), we estimated *C/τ* for many species grown in laboratory conditions, by mapping genome sequencing libraries on selected 100*kb* regions centered on the Ori and Ter loci, localized by compositional analysis [4]. Since growth conditions for genome sequencing are usually designed to achieve fast growth, we assume that these values are a proxy of each species’ maximum growth rate. Since only mero-oligoploids can divide faster than the time for chromosome replication (*C*), they can be identified as the fraction of species with *C/τ >* 1 i.e. *τ < C* (Figure 2b).

**Fig. 2.**
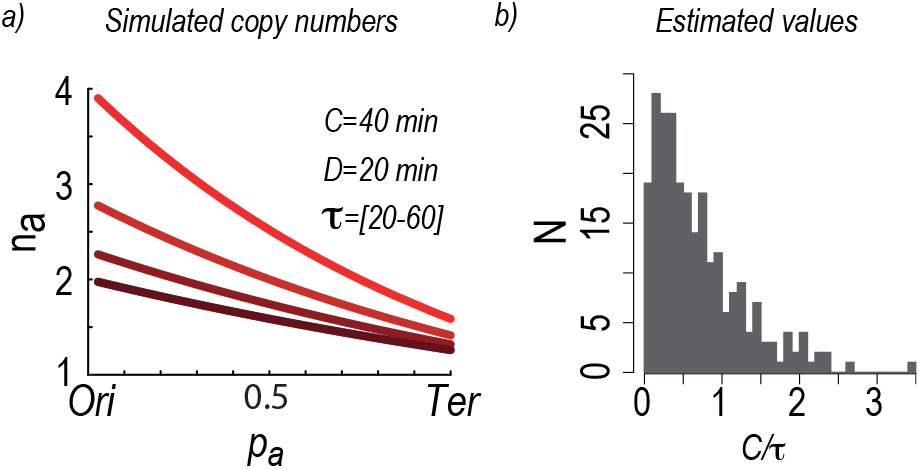
(a) Multiplicity *n*_*a*_ as a function of genomic position *p*_*a*_, computed in a range of 4 division times using Eq. 1; (b) Estimation of *C/τ* in many different species, using genome sequencing reads.

Since replication and transcription occur concomitantly in Prokaryotes, copy number variations directly affect the abundance of transcripts [1, 19, 18], therefore replication becomes an additional player in determining the transcriptome configuration of an organism.

In this work we integrate the above observations in a standard gene regulation model and we show that this approach highlights interesting evolutionary cues together with a deeper understanding of this global level of regulation and its interactions with transcriptional regulation.

## Results

### Mathematical formalization

A standard model describing the evolution in time of the abundance of a transcript *a*, under the regulation of a transcription factor *T* is:

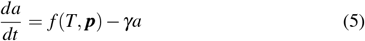

Where *γ* is the degradation rate, assumed to be constant across conditions and genes; *f* (*T*, ***p***) models the transcription rate of a promoter, as a function of the abundance of a transcriptional regulator *T* plus parameters. Often, this is a sigmoid (Eq. 6) or hyperbolic function that can be an increasing (activation) or decreasing (repression) with respect to the regulator’s abundance [7]:

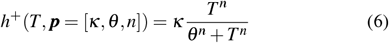

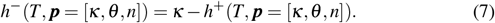

Above, *κ* is the maximum transcription rate achievable by this promoter at saturating regulator concentrations and *θ* (in units of concentration), is the regulator’s abundance inducing an half-maximal rate. Implicitly, this function represents the output from a single copy of the promoter. However, since prokaryotic genes can be transcribed just after their replication, it is natural to integrate a locus’ copy number as a multiplier of transcription rate in Eq. 5:

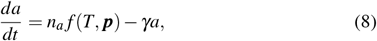

where *n*_*a*_ is the multiplicity of locus *a* defined in Eq. 1. Eq. 8 can be solved in *a* at the steady state (when 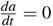):

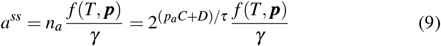

A solution that can be better discussed taking the log_2_:

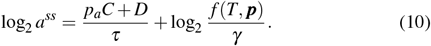

Eq. 10 highlights the different contributions to the steady state abundance of the transcript; as expected, the relative activity of the regulator with respect to the degradation rate is one, together with the dependence on division time and locus position. Therefore, changes in doubling time can affect the abundance of a transcript even with no change in the regulator’s activity. Since the two contributions can be considered as potentially independent, one pertinent question at this point is how sensitive the steady state transcript abundance is with respect to (i) the replication induced copy number variations and (ii) changes in the regulator’s activity.

### Sensitivity Analysis

To evaluate how the transcript steady state abundance depends on the two contributions in a quantitative way, we derived analytical formulas for the sensitivity of the solution to the two parameters of interest. Sensitivities can be calculated by partial derivation of the steady state expression with respect to one model parameter at time, obtaining:

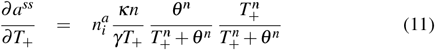

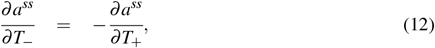

for the two possible regulatory effects, and:

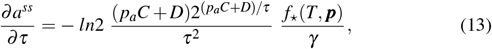

where *f* is the original regulatory function (positive or negative).

Sensitivities are represented in Figure 3 for biologically meaningful parameter ranges (*T* ∈ [0.1, 250] *µmol*, i.e. *≈* 11.3 log fold change around *θ* and *τ ∈* [30, 70]*min*). In Figure 3a we show the sensitivity of steady state transcript abundance to division time, when the transcription factor is constant, for all genomic positions. As expected, genes located closer to the origin are more sensitive to copy number changes; the negative sign comes out from the inverse relationship of division time and copy number i.e. when *τ* decreases (cells divide faster), a locus copy number grows. Additionally, copy number and sensitivity are positively correlated. In Figure 3b we show that the sensitivity to a positive regulator with a sigmoidal functional response acquires a positional pattern once replication is accounted for. As before, the sensitivity is higher nearby the origin and it increases at shorter division times (or equivalently, toward the origin). In Fig.3c the functional response is of the hyperbolic type; sensitivity in this case is maximum at very low regulator abundances and increases with copy number; the usual sigmoidal function instead translates into a log-normal shape, with a global maximum just before the regulator reaches a concentration *θ* (Fig. 3d). Increasing the steepness parameter (*n*) in the Hill function, makes the range of non-zero sensitivity narrower (not shown). The above clearly illustrates that copy number variations and genomic position affect the sensitivity to the regulator, therefore replication can introduce a positional and growth rate-dependent pattern in input/output relationships of gene regulatory networks.

**Fig. 3.**
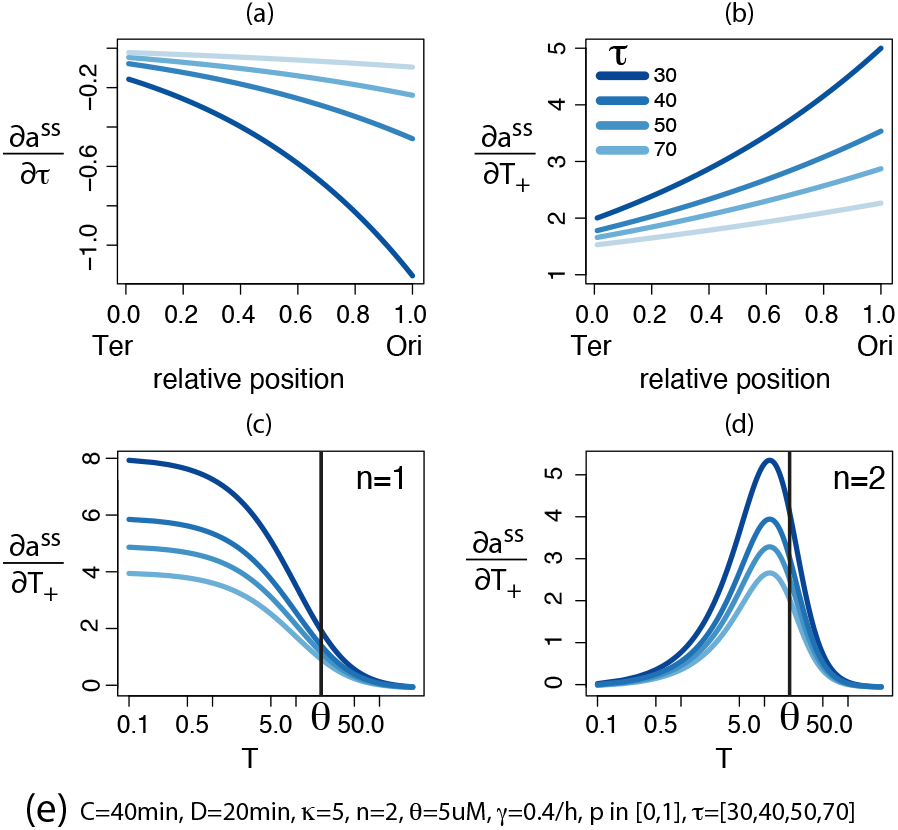
Sensitivity analysis. (a) Sensitivity to *τ* around different *τ* values, relative to genome position. (b) Sensitivity to transcription factor abundance (*T*) at various *τ* values (*T* = *const* = *θ*); (c) Sensitivity to transcription factor abundance at various *τ* values, relative to*T* itself for hyperbolic promoters; (d) as in (c) for a sigmoidal functional response; (e) parameters used, if not otherwise indicated.

### Variance decomposition

Since the steady state transcript level depends on both replication and transcriptional control, its variance across conditions must also be a function of both. Using Eq. 10 and the rule for calculating the variance of a sum of functions, we get:

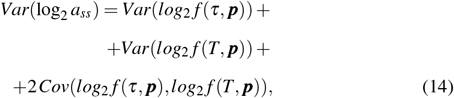

where *f* (*τ*, ***p***) = *n*_*a*_ as from Eq. 1, *Var*(*x*) is the variance of *x* and *Cov*(*x, y*) is the covariance of *x* and *y*. The variance and covariance *shape* depend on the family distribution of functions and the exact values of parameters, but general implications can be nonetheless derived. First, DNA replication introduces a position- and division time-dependent pattern to the variance: since genes at the origin have a wider range of copy numbers, their variance will be larger than for Ter-proximal genes. Second, if changes in copy number due to division time variations (*f* (*τ*, ***p***)) and the activity of the regulator (*f* (*T*, ***p***)) are uncorrelated, the covariance term in Eq. 14 will be zero. Therefore, the variance of the log-transformed transcript abundance is additive in those of the two functions. On the other hand, when the covariance term is positive, the variance of the transcript will increase faster than in the purely additive case. This can be interpreted as a replication-induced amplification of the activity of the regulator and could therefore be particularly advantageous for genes involved in growth-rate dependent processes (e.g. *σ* 70-dependent promoters). This reasoning provides a formal explanation to the documented tendency of *E. coli* and other fast-growing species to keep genes benefiting from high expression levels toward the origin [21]. On the converse, when the covariance term is negative, the result is a reduction of the variance with respect to the uncorrelated case (Figure 4).

**Fig. 4.**
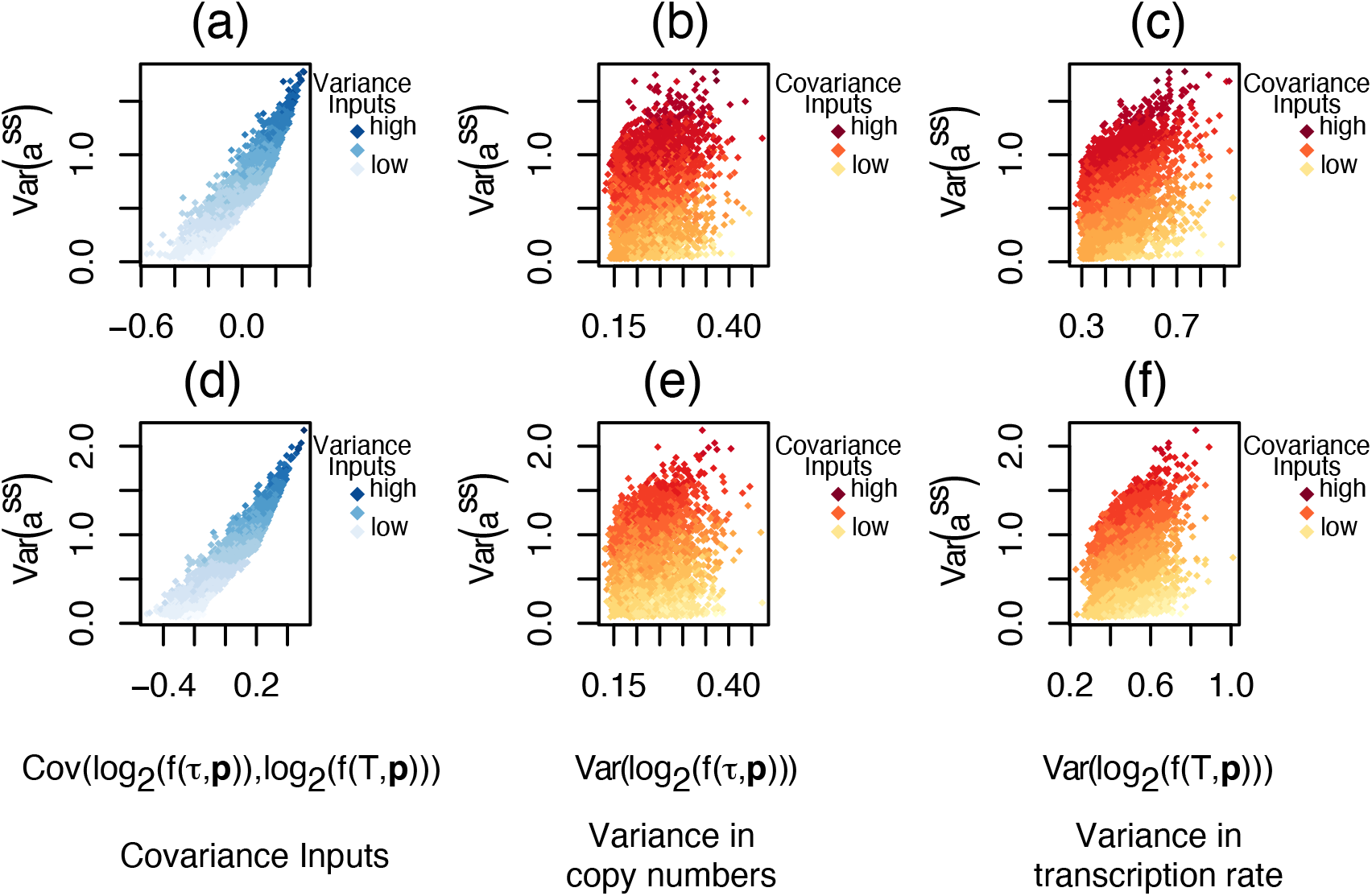
Variance - covariance relationships. *a,b* and *c* refer to the situation when *T* is an activator. *d, e, f* refer to *T* as a repressor. a,d) Relationship of the variance in the output with the covariance in the inputs. Data point colors corresponds to the sum of the variances in the inputs (light blue=low, blue=high); b, e) Relationship of the output with the variance in multiplicity. The color code corresponds to the covariance term (yellow=low, red=high); c, f) Relationship of the variance in the output with the variance in transcription rate. Color is as above. The variance in the output at steady state is strongly and positively correlated with the covariance term; the correlation with the inputs, albeit significant, are smaller.

We can therefore say that replication and transcription can cooperate or counteract each other, with a likely different impact on the Input/Output relationships of the system.

### Input/Output Analysis

To study how the covariation in the inputs (multiplicities, *f* (*τ*, ***p***) and transcription rates, *f* (*T*, ***p***)) might affect the Input/Output relationships in the system, we rely on the simulations described in methods. Shortly, by generating inputs at different covariance levels, we solve the system and check how the output (*a*^*ss*^) relates to the inputs.

Results of this analysis are summarized in Tables 1-4. We consider positive and negative regulation, and promoters with functional response of type II (hyperbolic) and III (sigmoidal). The emerging pattern can be summarized as: (i) positive correlation in the inputs **always** results in significant correlations of the output with both inputs. This should be a fairly common case for regulators not depending on post-translational activation, as their gene expression will be affected by division time changes in a way similar to their targets; (ii) Type III promoters tend to reflect the expected correlation with the regulator’s function but often not with multiplicity, basically disentangling the transcript’s abundance from the latter; (iii) Type II promoters tend to produce outputs that are instead more correlated with multiplicity than the regulator’s function; when *T* is a repressor it is more effective in defining patterns (ii) and (iii). (iv) However, the interference of copy numbers and regulator’s activity often masks the expected input/output relationship of *a*^*ss*^ and transcription rate. While being aware that the exact values of correlations and their significance depends on the parameters, by testing biologically meaningful regulator concentrations and division times, we highlight here that the interaction of copy number variations induced by changes in division time and transcription factor mediated regulation can result in an extended range of input/output relationships: positively correlated inputs provide a way to amplify the signal mediated by the regulator; on the converse, negative regulation on a type III promoter may provide a way to render a transcript’s abundance relatively independent from multiplicity.

**Table 1.**
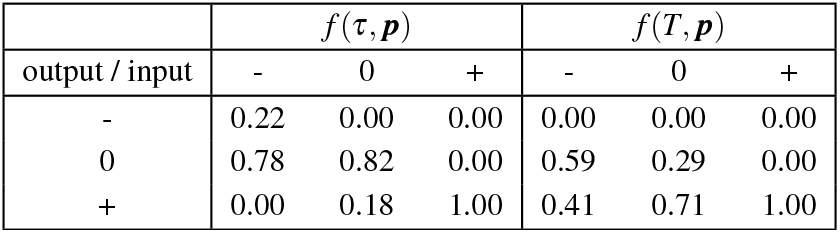
Input/output correlation analysis for an activator. (*κ* = 10, *n* = 2, other parameters are indicated in the main text). Rows corresponds to correlations of the output (*a*^*ss*^) with the function indicated in the header of each column block. In all cases, + (*−*) means a significant (*p ≤* 0.0001) positive (negative) correlation and 0 means the correlation is not significant. Columns concern the correlation in the inputs (*corr*(*f* (*τ*, ***p***), *f* (*T*, ***p***))). The “ − ” columns for instance mean that in the samples with inputs negatively correlated, *a*^*ss*^ is negatively correlated with multiplicity in 22% of the cases while the correlation is non significant in the remaining 78%; additionally, in only 41% of those cases, a significant and positive correlation with the transcription rate was detected; therefore, negatively correlated inputs often result in loss of correlation of the transcript abundance with respect to the regulator’s function. If the inputs are uncorrelated, the outputs also can become uncorrelated from the inputs. Differently, positively correlated inputs always result in high correlations in the outputs.

| output / input | $f(\tau, \mathbf{p})$ | | | $f(T, \mathbf{p})$ | | |
| --- | --- | --- | --- | --- | --- | --- |
|  | - | 0 | + | - | 0 | + |
| - | 0.22 | 0.00 | 0.00 | 0.00 | 0.00 | 0.00 |
| 0 | 0.78 | 0.82 | 0.00 | 0.59 | 0.29 | 0.00 |
| + | 0.00 | 0.18 | 1.00 | 0.41 | 0.71 | 1.00 |

**Table 2.** Same as Table 1 but with *n* = 1.

| output / input | $f(\tau, \mathbf{p})$ | | | $f(T, \mathbf{p})$ | | |
| --- | --- | --- | --- | --- | --- | --- |
|  | - | 0 | + | - | 0 | + |
| - | 0.00 | 0.00 | 0.00 | 0.22 | 0.00 | 0.00 |
| 0 | 0.39 | 0.25 | 0.00 | 0.78 | 0.72 | 0.00 |
| + | 0.61 | 0.75 | 1.00 | 0.00 | 0.28 | 1.00 |

**Table 3.** Same as Table 1, for a negative regulator. In this case, a negative correlation in the inputs translates in the loss of correlation of the output and multiplicity, while correlation with transcription rate is significant in almost 80% of the cases.

| output / input | $f(\tau, p)$ | | | $f(T, p)$ | | |
| --- | --- | --- | --- | --- | --- | --- |
|  | - | 0 | + | - | 0 | + |
| - | 0.00 | 0.00 | 0.00 | 0.00 | 0.00 | 0.00 |
| 0 | 1.00 | 0.83 | 0.00 | 0.21 | 0.25 | 0.00 |
| + | 0.00 | 0.17 | 1.00 | 0.79 | 0.75 | 1.00 |

**Table 4.** Same as Table 3 but with *n* = 1.

| output / input | $f(\tau, p)$ | | | $f(T, p)$ | | |
| --- | --- | --- | --- | --- | --- | --- |
|  | - | 0 | + | - | 0 | + |
| - | 0.00 | 0.00 | 0.00 | 0.00 | 0.00 | 0.00 |
| 0 | 0.07 | 0.30 | 0.00 | 1.00 | 0.78 | 0.00 |
| + | 0.93 | 0.70 | 1.00 | 0.00 | 0.22 | 1.00 |

## Conclusions

The relationship between genomic position and transcription rate due to replication-induced copy number variations introduces an additional layer of regulation for modulating gene expression. This hypothesis is congruent with papers demonstrating that gene position is a good predictor of gene expression levels [1, 2, 6, 16, 19, 20] and that there are genome organization patterns that can be explained consequently. In this work, we studied the interaction of replication-induced copy number variations and transcriptional regulation, from a theoretical perspective. By deriving analytical formulas for the sensitivity of steady state transcript’s abundance to both division time and regulator’s activity we showed that replication introduces a position- and division time-dependent pattern to the sensitivity of a transcript’s abundance with respect to the regulator’s activity. In this way, the cells have an additional mean to modulate the sensitivities of promoters on the basis of their position on the ori/ter axis. We highlight that the contribution of copy numbers and transcriptional regulation can interfere *constructively* or *destructively* in determining the variance of transcripts, and that this produces an enriched set of regulatory outputs that can be manipulated by controlling how the inputs are related to each other. A recent work [23] showed that constitutive promoters with the highest sensitivity to growth rate changes (after the copy number effect is removed), tend to be located toward the origin, and that their position is moreover conserved across many species. This could be explained by the evolutionary exploitation of the constructive effect of copy number and regulation variations highlighted by our treatment.

The decoupling of the variance of steady state transcript abundance into its components, shows that detecting target-regulator dependencies using gene expression data could be hampered by division time variations across conditions. Indeed, predicting the multiplicity of loci from gene expression data may enable to get rid of the variance introduced by replication, when the target is for instance promoter or regulator kinetic parameter estimation. Indeed, Eq. 10, highlights that even with constant regulator activity and degradation rate, growth in conditions affecting division times, would translate into different abundances at steady state; however, those fluctuations would be considered as genuine variations in regulator’s activity and incorporated into the parameters.

Similarly, when reconstructing gene regulatory networks using gene expression compendia (e.g. [8] replication can mask the dependency of targets with respect to the regulator; therefore, one may devise a correction of gene expression data to reduce the effect of copy numbers and consequently to highlight the relationship with regulator’s abundance.

We conclude adding that this work also suggests that multiplicity may help in the estimation of gene-specific dispersion factors in differential gene expression analysis (e.g. [14]) when replicates are in small number, and the conditions differ in terms of division times.

## Methods

Scripts in R for reproducing all the analysis and figures in the paper are available at the team’s repository on github: Comparative Systems Biology Lab.

### Simulations for Covariance and I/O analysis

Simulations corresponding to data points in Figure 4 and Tables 1-4 were done with each iterations consisting of the following.

1. Copy numbers and transcription factor abundances were generated to have a known covariance using mvrnorm with *µ* = [0, 0] and variance-covariance matrix:

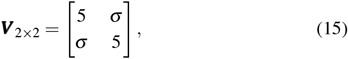

where *σ* is the covariance and the diagonal entries are variances. We tested 50 covariance levels from *−*5 to 5 in 0.2 steps; each covariance level was tested 20 times.
2. Scale values to have *τ ∈* [20, 70]*min* and *T ∈* [2, 12]*µmol*.
3. Calculate steady state transcript abundances for the above parameters using Eq. 9 and the following values *κ* = 10, *θ* = 5*µmol,C* = 40*min, D* = 20*min, γ* = 0.0067*min*^*−*1^. Several parameter combinations were tested but the results are not significantly different.
4. Variance/Covariance and Correlations:
  - Calculate variances and covariances (Eq. 14), shown in Figure 4.
  - Calculate Input/Output and Input/Input correlations for Tables 1-4.

## Author contributions statement

MB conceived the idea. MB and PG developed the models and performed the analysis, students participated actively to the discussion.

## Acknowledgments

MB wish to thank the students of his 2022 course at the Quantitative Biology Master Degree of the University of Milan for participating to the development of this idea; they were: Mario Deligios, Daniele di Bella, Greta Carola Giannini, Elahe Hassanbeiki, Frederik Espersen Knudsen and Alberto Peruzzi.

**Matteo Brilli, Ph.D**. is an Associate Professor in Microbiology at the University of Milan (Italy). He is interested in basic research and general questions that he approaches by integrating his variegated expertise in bioinformatics and systems biology.

**Pietro Gemo.** At the time of writing he was an undergraduate of the Master in Quantitative Biology of the University of Milan and now a Ph.D. candidate at the Max Planck Institute for Infection Biology (Berlin).

